# Inside-out regulation of E-cadherin conformation and adhesion

**DOI:** 10.1101/2020.05.02.074187

**Authors:** Ramesh Koirala, Andrew Vae Priest, Chi-Fu Yen, Joleen S. Cheah, Willem-Jan Pannekoek, Martijn Gloerich, Soichiro Yamada, Sanjeevi Sivasankar

**Author notes:** Equal contribution (Ramesh Koirala, Andrew Vae Priest).

## Abstract

Cadherin cell-cell adhesion proteins play key roles in tissue morphogenesis and wound healing. Cadherin ectodomains bind in two conformations, X-dimers and strand-swap dimers, with different adhesive properties. However, the mechanisms by which cells regulate ectodomain conformation are unknown. Cadherin intracellular regions associate with several actin-binding proteins including vinculin, which are believed to tune cell-cell adhesion by remodeling the actin cytoskeleton. Here, we show at the single molecule level, that vinculin association with the cadherin cytoplasmic region allosterically converts weak X-dimers into strong strand-swap dimers, and that this process is mediated by myosin II dependent changes in cytoskeletal tension. We also show that in epithelial cells, ∼70% of apical cadherins exist as strand-swap dimers while the remaining form X-dimers, providing two cadherin pools with different adhesive properties. Our results demonstrate, for the first time, the inside-out regulation of cadherin conformation and establish a mechanistic role for vinculin in this process.

**SIGNIFICANCE STATEMENT:** Cadherin cell-cell adhesion proteins play key roles in the formation and maintenance of tissues. Their adhesion is carefully regulated to orchestrate complex movement of cells. While cadherin ectodomains bind in two conformations with different adhesive properties, the mechanisms by which cells regulate the conformation (and consequently adhesion) of individual cadherins are unknown. Here, we demonstrate that the association of intracellular vinculin to the cadherin cytoplasmic region, regulates cadherin adhesion by switching ectodomains from a weak binding to the strongly adhesive conformation. In contrast with the prevailing view which suggests that vinculin regulates adhesion solely by remodeling the cytoskeleton, we show that vinculin can directly modulate single cadherin ectodomain conformation and that this process is mediated by changes in cytoskeletal tension.

## INTRODUCTION

E-cadherins (Ecads) are essential, calcium dependent cell-cell adhesion proteins that play key roles in the formation of epithelial tissue and in the maintenance of tissue integrity. Ecad adhesion is highly plastic and carefully regulated to orchestrate complex movement of epithelial cells and dysregulation of adhesion is a hallmark of numerous cancers ^1^. However, little is known about how cells dynamically regulate the biophysical properties of individual Ecads.

The extracellular region of Ecads from opposing cells bind in two distinct *trans* orientations: strand-swap dimers and X-dimers (Figure 1A-B). Strand-swap dimers are the stronger cadherin adhesive conformation and are formed by the exchange of conserved Tryptophan (Trp) residues between the outermost domains of opposing Ecads ^2–4^. In contrast, X-dimers, which are formed by extensive surface interactions between opposing Ecads, are a weaker adhesive structure and serve as an intermediate during the formation and rupture of strand swap dimers ^5–7^. Using cell-free, single molecule experiments, we previously showed that X-dimers and strand-swap dimers can be distinguished based on their distinctly different response to mechanical force. When a strand-swap dimer is pulled, its lifetime decreases with increasing force, resulting in the formation of a slip bond ^8, 9^ (Figure 1B). In contrast, an X-dimer responds to pulling force by forming a catch bond, where bond lifetime initially increases up to a threshold force, and then subsequently decreases ^8, 10^ (Figure 1B). It has also been shown that wild type Ecad ectodomains in solution can interconvert between X-dimer and strand-swap dimer conformations ^9, 11^. However, the biophysical mechanisms by which Ecad conformations (and adhesion) are regulated on the cell surface are unknown.

**Figure 1.**
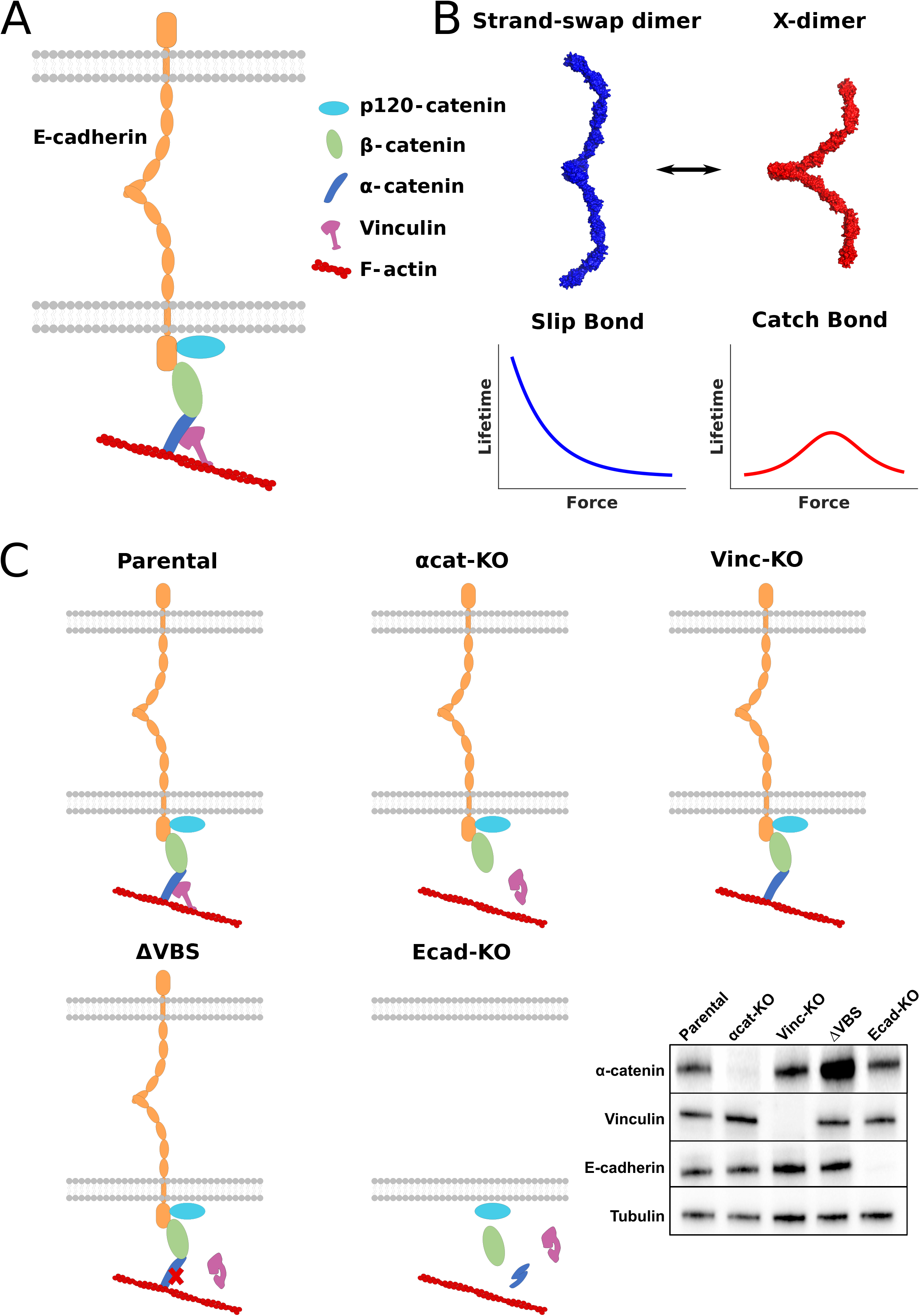
Overview of experiment. **(A)** The extracellular region of Ecad from opposing cells mediate adhesion. The cytoplasmic region of Ecad associates either directly or indirectly with p120 catenin, β-catenin, α-catenin, vinculin, and F-actin. **(B)** Strand-swap dimers form slip bonds (blue) while X-dimers form catch bonds (red). Ecads interconvert between these two dimer conformations. Structures were generated from the crystal structure of mouse Ecad (PDB: 3Q2V); the X-dimer was formed by alignment to an X-dimer crystal structure (PDB: 3LNH). **(C)** Graphics showing the cell lines used in experiments and western blot analysis of corresponding cell lysates.

The cytoplasmic region of Ecad associates with the catenin family of proteins: namely p120-catenin, β-catenin and α-catenin. The Ecad-catenin complex, in turn, links to filamentous actin (F-actin) either by the direct binding of α-catenin and F-actin or by the indirect association of α-catenin and F-actin via vinculin ^12^ (Figure 1A). Adhesive forces transmitted across intercellular junctions by Ecad induce conformational changes in α-catenin ^13, 14^, strengthen F-actin binding ^15^ and recruit vinculin to the sites of force application ^16, 17^. However, vinculin and α-catenin do not merely serve as passive cytoskeletal linkers; they also dynamically modulate cytoskeletal rearrangement and recruit myosin to cell-cell junctions ^13, 18–20^. Studies show that α-catenin and vinculin play important roles in strengthening and stabilizing Ecad adhesion: bead twisting experiments show force-induced stiffening of Ecad-based junctions and cell doublet stretching experiments demonstrate reinforcement of cell-cell adhesion in vinculin and α-catenin dependent manners ^18, 19, 21^.

Currently, actin anchorage and cytoskeletal remodeling are assumed to be the exclusive mechanisms by which α-catenin and vinculin strengthen Ecad adhesion ^22–24^. Here, we directly map the allosteric effects of cytoplasmic proteins on Ecad ectodomain conformation and demonstrate, at the single molecule level, that vinculin association with the Ecad cytoplasmic region switches X-dimers to strand-swap dimers. We show that cytoskeletal tension, due to vinculin mediated recruitment of myosin II, regulates Ecad ectodomain structure and adhesion. Finally, we demonstrate that only ∼50% of Ecads are linked to the underlying cytoskeleton and that while about 70% of Ecads form strand-swap dimers, the remaining form X-dimers, which provides cells with two Ecad pools with different adhesive properties.

## RESULTS

### Single molecule Atomic Force Microscope (AFM) measurements of specific Ecad *trans* adhesion

We measured interactions between recombinant Ecad extracellular regions immobilized on AFM cantilevers and Ecad endogenously expressed on the apical surface of confluent Madin-Darby Canine Kidney (MDCK) cell monolayers (Figure 2A). To ensure that we measured single Ecad unbinding events, we immobilized Ecad on the AFM tip at low density and pressed on the cell surface with a low initial contact force (mean pressing force = 59 pN; Figure S1). Poisson statistics predict that even the maximum event rate we measure (19%; Figure 2B) corresponds to ∼90% probability of measuring single molecule events.

**Figure 2.**
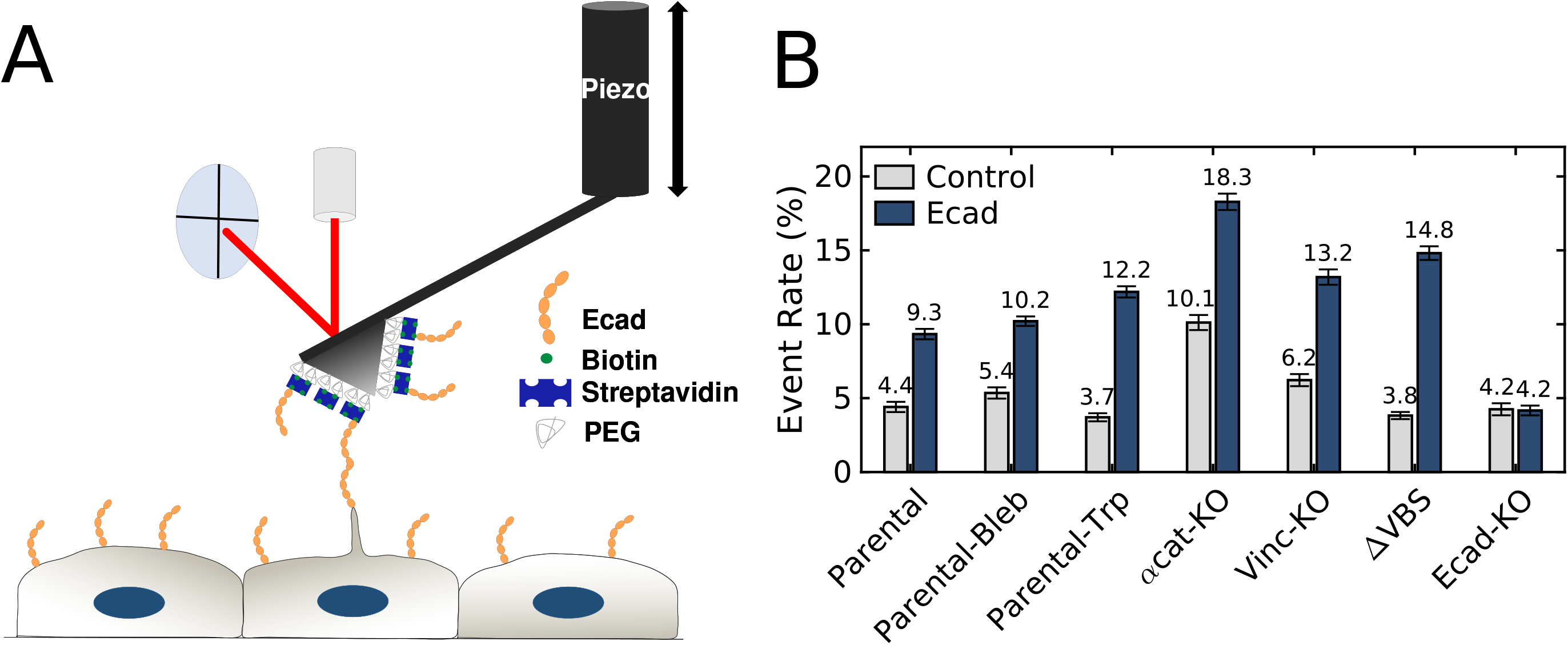
Measuring specific and non-specific interaction using AFM. **(A)** Biotinylated Ecad monomers were immobilized on AFM cantilevers functionalized with Polyethylene glycol (PEG) and Streptavidin as described in the methods section. Interactions between Ecads on the cantilever and Ecads expressed on the apical surface of MDCK cells were measured. **(B)** Binding probabilities were measured using AFM cantilevers lacking Ecad (control experiment, light gray) or AFM cantilevers decorated with Ecad (Ecad, dark blue). Total number of Ecad/control measurements performed on each cell line were 6793/3361 measurements on parental cells, 8593/3719 measurements on parental-Bleb, 7268/4858 measurements on parental-Trp, 4291/3666 measurements on vinc-KO cells, 4518/3381 measurements on αcat-KO cells, 6124/6070 measurements on ΔVBS cells, and 3672/2449 measurements on Ecad-KO cells. Error bars are bootstrapped standard deviations.

Our experiments were performed with the following MDCK cells (Figure 1C): parental cells (*parental*), vinculin knockout cells (*vinc-KO*), α-catenin knockout cells (*αcat-KO*) ^25^, αcat-KO cells rescued with α-catenin lacking the vinculin binding site (*ΔVBS*) ^26^, and Ecad knockout cells (*Ecad-KO*). We also performed measurements with parental cells in the presence of free tryptophan (*parental-Trp*) in order to trap Ecad in an X-dimer conformation ^8^. Additionally, we performed experiments in the presence of blebbistatin (*parental-Bleb*) in order to reduce myosin II dependent cytoskeletal contractility. We quantified the specific and nonspecific binding rates for every cell line by measuring single unbinding events using either AFM tips that were decorated with Ecad (specific binding) or AFM tips that lacked Ecad (nonspecific binding). All cell lines, except Ecad-KO, showed a significant increase in single unbinding events when the AFM tips were decorated with Ecad (Figure 2B). In contrast, in the Ecad-KO cells, event rates using AFM tips with/without Ecad were similar to each other and were comparable to the nonspecific event rates measured with the other cell lines (Figure 2B). Taken together, this confirmed that the measured specific events corresponded to Ecad-Ecad binding interactions.

Ecad conformation and cytoskeletal-linkage in each cell line were determined from previously reported ‘signatures’ in the measured force curves. Since force measurements have shown that pulling on a cell surface protein that is not linked to the cytoskeleton results in the formation of a *membrane-tether* ^27, 28^ (Figure 3A), we used membrane-tethers to identify Ecad that are uncoupled from the actin cytoskeleton (Figure 3B). Previous studies have also shown that pulling on a transmembrane protein that is firmly bound to the underlying cytoskeleton results in a linearly elastic response to pulling force ^29, 30^ (Figure 3C). Since these *jump* events occur when interactions between proteins on the AFM tip and cell surface are weaker than the protein-cytoskeleton linkage, we interpreted jumps as a signature of actin-linked Ecads that form weak *trans* dimers, which rupture before failure of the Ecad-cytoskeletal linkage (Figure 3D). Finally, if the Ecad *trans* dimers adhere robustly compared to the Ecad-cytoskeletal linkage, an initial steep increase in force is measured which relaxes viscoelastically when the Ecad-cytoskeletal bond ruptures, resulting in the formation of a *cyto-tether* (Figure 3E-F). To further validate tether formation, we fit the membrane-tether and cyto-tether force-extension curves to the standard linear solid (SLS) model (SI Note 1) ^31^. The SLS fitting parameters we obtained for the tethers, were consistent with previous values obtained for tether formation on Jurkat cells and T-cells (Table S1) ^31, 32^.

**Figure 3.**
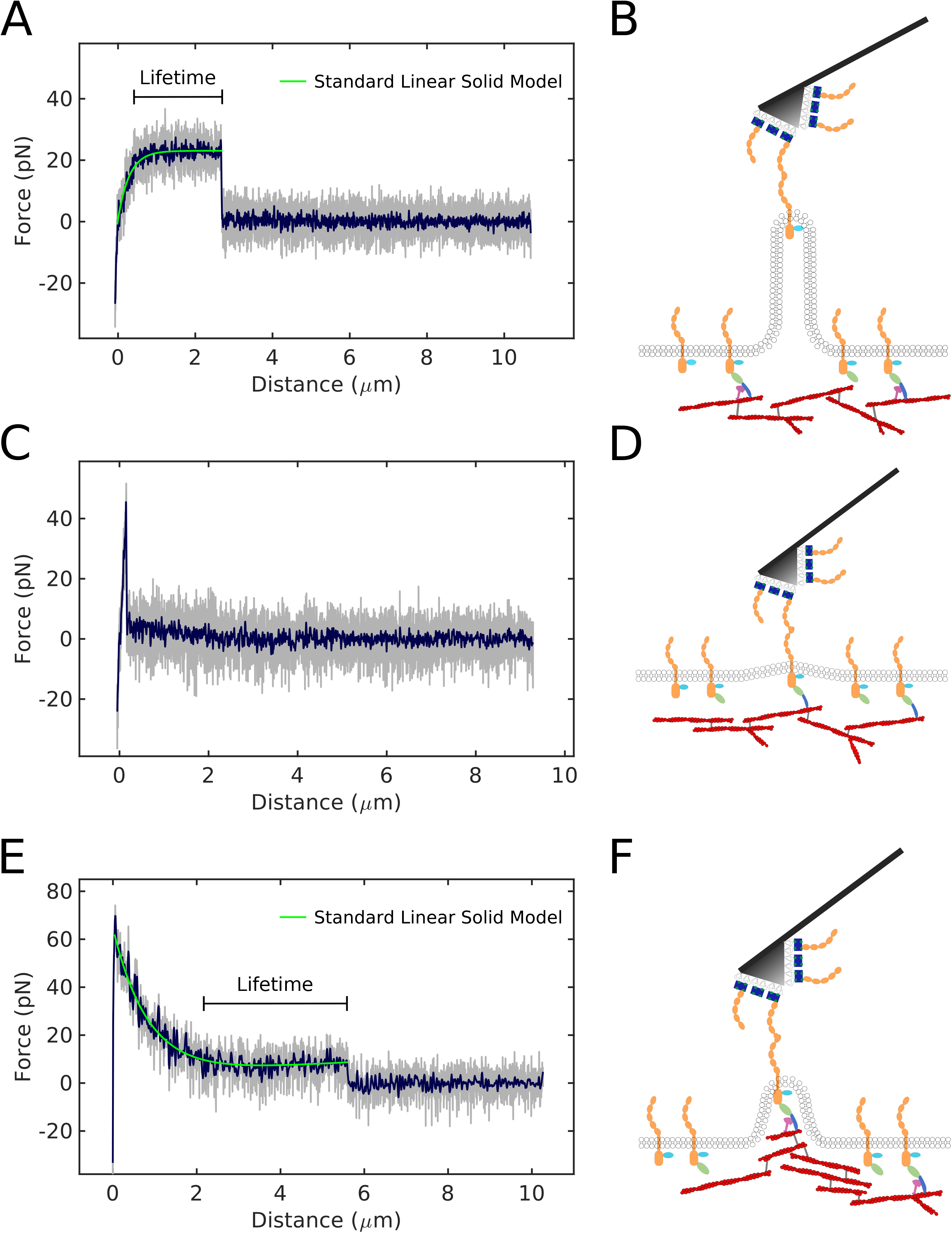
Force ‘signatures’ used to identify cytoskeletal coupling and strength of *trans* adhesion. **(A)** Typical force curve showing a membrane-tether which extends at a constant force. **(B)** Membrane-tethers are measured when an Ecad, which is not linked to the actin cytoskeleton is pulled. **(C)** Jump events exhibit a linear increase in force and are measured upon pulling **(D)** an Ecad *trans* dimer that is weaker than its cytoskeleton coupling. **(E)** Typical force curve showing a cyto-tether with an initial increase in force that viscoelastically relaxes as a membrane-tether is formed. **(F)** Cyto-tethers are measured when a robust Ecad *trans* dimer that is linked to the actin cytoskeleton is pulled. Raw data (gray) was acquired at 25 kHz and smoothed (blue) to 111 Hz. Green curve is the Standard Linear Solid model fit to membrane-tethers and cyto-tethers. Black bars indicate bond lifetimes determined from membrane-tethers and cyto-tethers.

### About 50% of non-junctional Ecads are coupled to the actin cytoskeleton

We first compared the fraction of Ecad uncoupled from the cytoskeleton (membrane-tethers) to the fraction of Ecads that are coupled to cortical actin (cyto-tethers and jumps). Our data showed that in parental cells, 55% ± 9% of Ecad on the apical surface of the cell are uncoupled from the actin cytoskeleton (Figure 4A). Since α-catenin is believed to be vital in mediating Ecad linkage to F-actin, we next measured the cytoskeletal linkage of Ecad in αcat-KO cells. As anticipated from previous biochemical and cell biological results ^33, 34^, our data showed that the vast majority of Ecad (92% ± 12%) in αcat-KO cells are uncoupled from the actin cytoskeleton (Figure 4A). In contrast, the fraction of uncoupled Ecads in both vinc-KO and ΔVBS cells, where Ecads are presumably weakly linked to F-actin by α-catenin, but where this linkage is not reinforced by vinculin ^13, 35^, was comparable to parental cells (65% ± 9% for vinc-KO and 45% ± 4% for ΔVBS cells; Figure 4A). Taken together, this data confirms the obligatory role of α-catenin in mediating Ecad linkage to F-actin. These direct, single molecule measurements of Ecad-actin coupling on the apical surface of live cells are in excellent agreement with previous single particle tracking of Ecad on the cell surface that inferred approximately 50% of Ecads on the apical surface of cells are coupled to the actin cytoskeleton ^36^.

**Figure 4.**
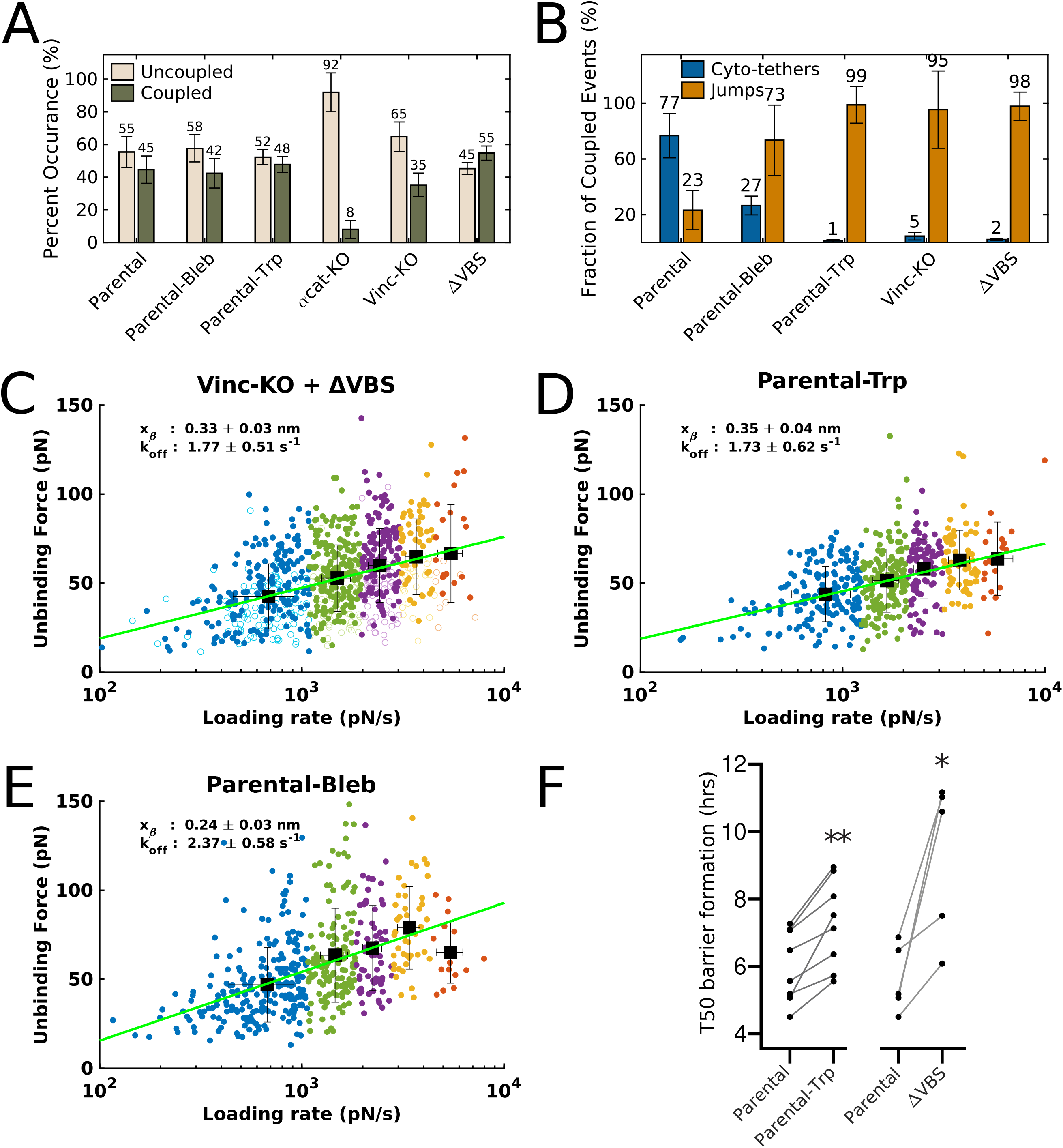
Role of α-catenin and vinculin in Ecad-cytoskeleton linkage and adhesion strength. **(A)** Fraction of uncoupled (membrane-tether) and coupled (cyto-tether + jump) events. Fractions were calculated after subtracting the binding probability of the control experiment for each cell line. **(B)** Fractions of strong (cyto-tether) and weak (jump) Ecad *trans* dimer events. Error bars are the propagated errors obtained from bootstrapped standard deviations. Dynamic force spectroscopy of jump events in **(C)** vinc-KO + ΔVBS, **(D)** parental-Trp, and **(E)** parental-Bleb cells. Loading rates and peak forces in jump events were bootstrapped 1000 times then clustered using k-means and fit to the Bell-Evans model (green line) using weighted nonlinear least-square fitting. Data points represent single unbinding events, and colors correspond to data points from the same k-means cluster. The data from ΔVBS (filled circles) and vinc-KO (circles) cells in (C) were combined because the number of data points were insufficient for a statistically reliable calculation. Intrinsic off rate (k_off_) and distance to the transition energy barrier (x_β_) were obtained from the mean of the fitted parameters acquired by fitting the k-means clusters that were calculated from the bootstrapped samples. Errors in the fitted parameters correspond to standard deviations of the bootstrapped means. **(F)** Trans-epithelial resistance measurement of parental cells in the absence (Parental; n = 9) and presence (Parental-Trp; n = 8) of 2 mM tryptophan, or ΔVBS cells (ΔVBS; n = 5), following addition of 2 mM CaCl_2_ to induce Ecad dependent formation of the epithelial barrier. Barrier formation half times (T50) were calculated as the time-point at which the impedance signal surpassed 50% of the ultimate maximum impedance. * p = 0.0025, ** p = 0.0011; Paired t-test.

### Absence of vinculin binding weakens adhesion of individual Ecad

We next compared the fraction of weak (jumps) and strong (cyto-tethers) *trans* dimers, formed by cytoskeleton-coupled Ecad. Our data showed that in parental cells that contain both α-catenin and vinculin, 77% ± 16% of cytoskeleton-coupled Ecad bind robustly while the remaining 23% ± 14% of Ecad interact weakly (Figure 4B). In contrast, in vinc-KO cells, only 5% ± 3% of cytoskeleton-coupled Ecad bind strongly (Figure 4B). Similarly, in ΔVBS cells where vinculin cannot bind to α-catenin, only 2% ± 1% of cytoskeleton-linked Ecad bind in a robust binding conformation (Figure 4B). Taken together, this indicates that vinculin association with the Ecad cytoplasmic region is required for robust Ecad *trans* dimerization. It is important to point out that the strength of *trans* dimer adhesion in the absence of α-catenin could not be determined by comparing the fraction of cyto-tethers and jumps because these events were extremely rare in the αcat-KO cells.

Since an Ecad *trans* dimer can exist in two different conformations with different adhesive strengths — weaker X-dimer and stronger strand-swap dimer — we asked whether the weak (jumps) and strong (cyto-tethers) adhesive states of cytoskeleton-coupled Ecad represent different Ecad conformations. Since we have previously shown that Ecad ectodomains can be trapped in an X-dimer conformation simply by incubating them in free Trp ^8^, we performed AFM measurements on parental-Trp cells to identify the signature of X-dimer formed by cytoskeleton-coupled Ecad. Interestingly, almost all (99% ± 13%) cytoskeleton-bound Ecads in parental-Trp formed jump events (Figure 4B), suggesting that the jump events represent the Ecads coupled to the cytoskeleton that form X-dimers. This, along with exclusive presence of jumps in vinc-KO and ΔVBS (Figure 4B), indicates that the weakening of Ecad adhesion in the absence of vinculin binding is consistent with Ecad being trapped in an X-dimer conformation. Importantly, switching ectodomains into an X-dimer conformation did not decouple Ecad from the cytoskeleton and the fraction of cytoskeleton-coupled Ecad in the Trp treated cells (52% ± 5%) remained the same as parental cells (Figure 4A).

We also inferred the *trans* dimer binding conformation of cytoskeleton-coupled Ecad in the parental-Trp, vinc-KO and ΔVBS cells using dynamic force spectroscopy analysis (DFS) ^37^. Since the number of jump events in vinc-KO cells were insufficient to obtain statistically reliable results, we combined the vinc-KO jumps with the jumps measured with ΔVBS cells. The measured k_off_ for jump events for vinc-KO and ΔVBS (k_off_ = 1.77 ± 0.51 s^-1^; Figure 4C) was comparable to parental-Trp X-dimers (k_off_ = 1.73 ± 0.62 s^-1^; Figure 4D), and higher than previously published values for strand-swap dimers ^38–41^. Since a higher k_off_ corresponds to a weaker binding, these data further suggest that in the absence of vinculin association, cytoskeleton-coupled Ecad adopts a weaker X-dimer conformation.

Next, since cadherins trapped in an X-dimer conformation have delayed junction formation ^42^, we monitored the formation of cell-cell barriers in parental-Trp and ΔVBS cells using Electric Cell-Substrate Impedance Sensing (ECIS), in order to determine if the junction formation in these cell lines are consistent with Ecad adopting an X-dimer conformation. Our experiments measuring the barrier formation half time (T50), showed a significant delay in calcium-induced barrier formation in parental-Trp and ΔVBS cells compared to parental cells (parental-Trp 1.23 ± 0.65 hr delay in T50, ΔVBS 3.66 ± 2.26 hr delay in T50; Figure 4F). Importantly, delayed barrier formation with ΔVBS was consistent with previous reports ^26^. Taken together, our ECIS results indicate that both Ecad conformation and vinculin binding to α-catenin are important for the formation of cadherin junctions, and further suggest that vinculin is responsible for regulating Ecad conformation.

### Ecad ectodomain conformation is regulated by vinculin

Next, we proceeded to directly measure the binding conformation of Ecad ectodomains in the different cell lines by measuring if the Ecads on the cell surface form catch bonds (X-dimers) or slip bonds (strand-swap dimers). We measured the force *vs.* lifetimes of Ecad-Ecad bonds, using a method described previously ^43^. Briefly, since the force required to rupture membrane-tethers and cyto-tethers scales linearly with pulling velocity ^28, 44^ (Figure S2), and because these tethers extend at a constant force (after any initial viscoelastic relaxation) when pulled at a constant velocity ^27, 31^ (Figure 3A, 3E), we determined bond lifetimes at a range of forces from the persistence time of membrane-tethers and cyto-tethers.

We first measured the force *vs.* lifetime of Ecad endogenously expressed on the apical surface of parental cells. Fits of our data to the composite of a slip bond ^45^ and catch bond ^46^ model showed that 69% of Ecad in the parental cells form strand-swap dimers while the remaining 31% of Ecad occupy an X-dimer conformation (Figure 5A; Methods; Table S2). Upon Trp incubation, however, Ecads in parental cells switched their conformation and formed catch bonds characteristic of an X-dimer (Figure 5B).

**Figure 5.**
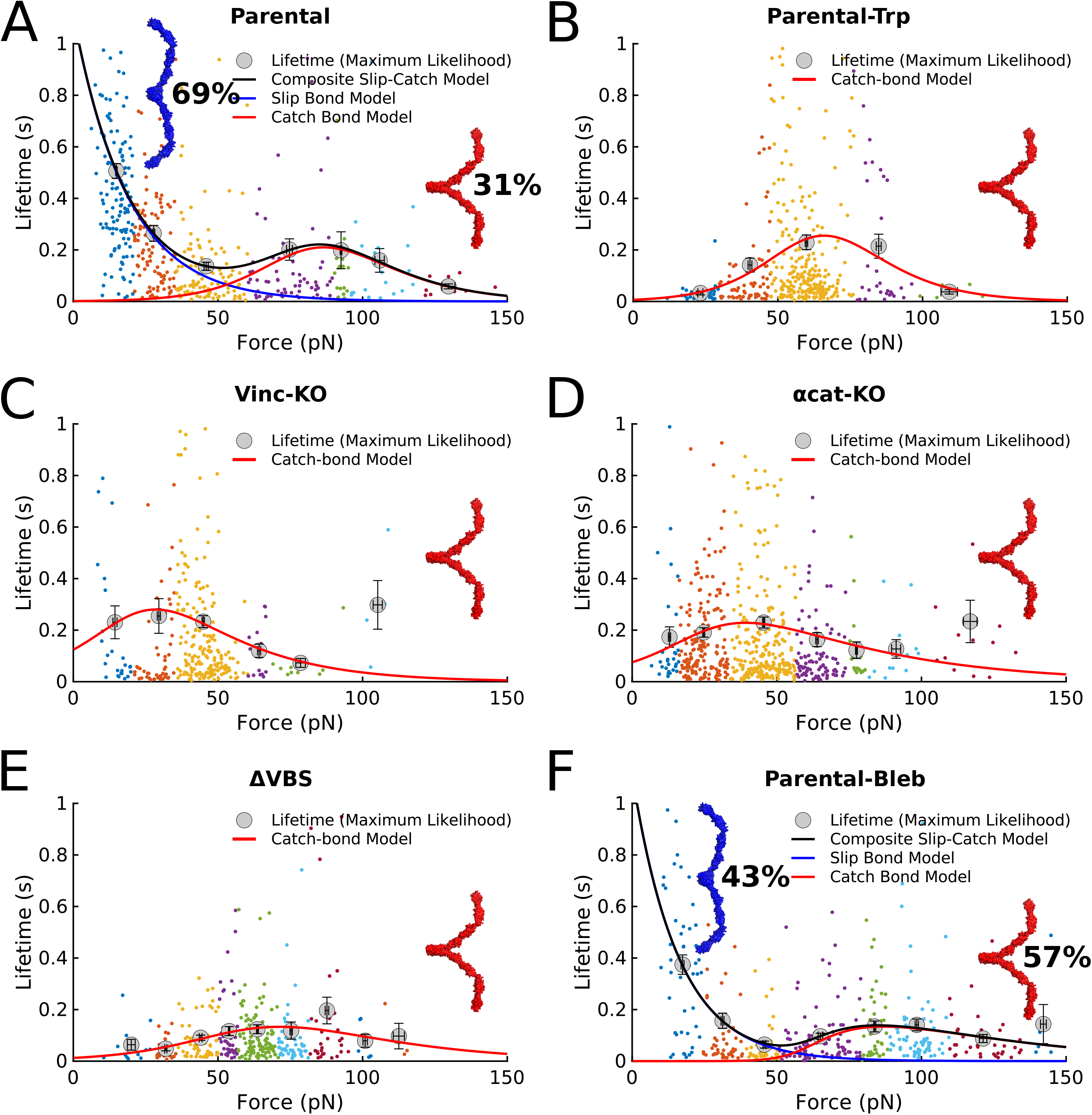
Measuring conformation of Ecad *trans* dimers using an AFM. Force *vs* lifetime profile for **(A)** parental cells show that 69% of Ecad form slip bonds (strand-swap dimer structure shown in blue) while the remaining 31% form catch bonds (X-dimer structure shown in red). **(B)** Parental-Trp, **(C)** vinc-KO, **(D)** αcat-KO, and **(E)** ΔVBS cells form catch bonds. **(F)** Inhibition of Myosin II using Blebbistatin reduces the fraction of slip bond forming Ecad. Parental-Bleb shows that 43% of Ecad form slip bonds while the remaining 57% form catch bonds. Solid lines represent fits to the corresponding models. Data was binned using a Gaussian mixture model and the lifetimes were determined using maximum likelihood estimation. The colored data points correspond to the data points in each respective bin. The data was fit by χ^2^ minimization and errors were obtained by bootstrapping. The y-axes were limited to 1s for clarity.

Since the bond-lifetimes for parental cells (Figure 5A) were obtained from all the tethers (a mix of cyto-tethers and membrane-tethers), we separately analyzed the force-dependent lifetimes of cyto-tether and membrane-tether events (Figure S3A, B). Consistent with cyto-tethers corresponding to strong Ecad-Ecad bonds, our analysis showed that cyto-tethers (which correspond to Ecad initially coupled to the cytoskeleton) exclusively formed slip-bonds. Ecad that were decoupled from the cytoskeleton (membrane-tether events) formed X-dimers and strand-swap dimers with almost equal probabilities (45% catch bonds: 55% slip bonds) (Figure S3A, B). This provided further evidence that the association of vinculin with the Ecad cytoplasmic domain is required for ectodomains to efficiently form strand-swap dimers.

As anticipated from our DFS experiments, Ecads in the vinc-KO cells formed catch bonds characteristic of an X-dimer conformation (Figure 5C) suggesting that vinculin association is necessary for Ecads to form robust strand-swap dimers. Similarly, Ecads in αcat-KO cells, where vinculin cannot associate with the Ecad cytoplasmic region ^12^, also formed catch bonds (Figure 5D). Finally, eliminating vinculin binding to α-catenin in ΔVBS cells also resulted in catch bonds (Figure 5E), confirming that association of vinculin with the Ecad cytoplasmic region via α-catenin is obligatory for transitioning from a weak X-dimer conformation to a robust strand-swap dimer conformation.

### Regulation of Ecad ectodomain conformation is myosin II dependent

Having identified vinculin as a key protein involved in allosterically regulating Ecad ectodomain conformation, we proceeded to identify the mechanism by which vinculin regulates Ecad structure. Since vinculin localizes to force bearing sites in the cell-cell junction and mediates the recruitment of myosin II ^18^, we hypothesized that Ecad ectodomain conformation could be regulated by myosin II mediated cytoskeletal contractile force. We therefore tested the effect of myosin II mediated force on Ecad conformation in parental cells by reducing cytoskeleton contractility using the myosin II inhibitor, blebbistatin (*parental-Bleb*). Strikingly, in parental-Bleb cells, the fraction of Ecad in strand-swap dimer conformation (43%; Figure 5F) decreased compared to parental cells (69% strand-swap dimer; Figure 5A). This indicated that myosin II dependent cytoskeletal contractility plays a role in driving strand-swap dimerization of Ecad ectodomains.

Additionally, unlike parental cells where the majority of cytoskeleton-coupled Ecad existed in the robust binding conformation (cyto-tethers), most of the cytosekeleton-coupled Ecad in parental-Bleb formed weak binding structures (jumps) (27% ± 7% cyto-tethers, 73% ± 25% jumps; Figure 4B). Importantly, the fraction of cytoskeleton-coupled Ecad in parental-Bleb (42% ± 9%; Figure 4A) remained the same as in parental cells. Finally, the measured k_off_ for jump events in parental-Bleb (2.37 ± 0.58 s^-1^; Figure 4E) was comparable to the values measured for X-dimers formed by parental-Trp, vinc-KO and ΔVBS cells (Figure 4C-D). Taken together, these data confirm that reduced myosin II dependent cytoskeletal contractility reduces the ability of Ecad to form robust strand-swap dimers.

Since forces generated by the actomyosin cytoskeleton can transmit to the Ecad cytoplasmic domain ^47^, we used steered molecular dynamics (SMD) simulations to test if force could convert X-dimers to strand-swap dimers (Figure 6A-B). We modeled cytoskeletal forces by pulling on the C-terminus of the EC1-2 X-dimer crystal structure (PDB ID: 3LNH) at five angles (0°, ±30°, ±40°), chosen to account for different orientations of actin filaments (Figure 6B). Our simulations showed that regardless of the pulling angle, all X-dimers converted to strand-swap dimers after a period of force application (Figure 6B-C). Root mean squared deviations (RMSDs) of the SMD structures, calculated relative to the strand-swap dimer crystal structure (PDB ID: 2QVF), converged to an average of 0.67 ± 0.12 nm confirming that the structures closely resembled strand-swap dimers (Figure 6C). These simulations confirm that when Ecad dimers are firmly coupled to the actin cytoskeleton by α-catenin and vinculin, myosin-II dependent force can convert X-dimers to strand-swap dimers.

**Figure 6.**
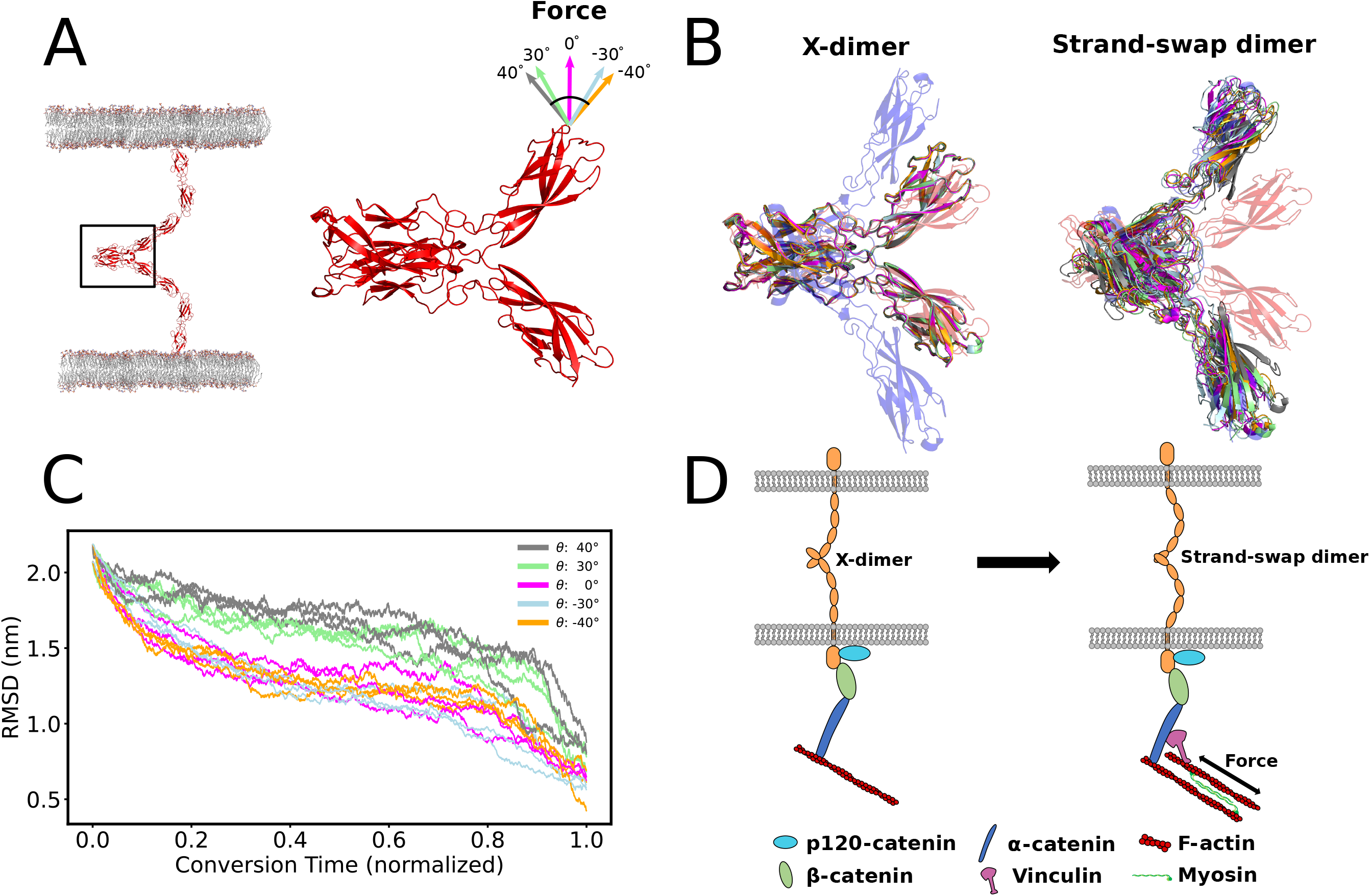
Cytoskeletal tension converts X-dimers to strand-swap dimers. **(A)** X-dimer formed by Ecad on opposing cells (left). X-dimer crystal structure (PDB ID: 3LNH) used in SMD simulations (right). The pulling directions are indicated by colored arrows. The full X-dimer (left) was simulated by aligning 3Q2V to 3LNH. **(B)** X-dimer conversion to strand-swap dimer. Simulated structures remain in an X-dimer conformation after stabilizing MD simulations (left). X-dimers become strand-swap dimers after pulling on the C-terminal of one Ecad at a range of angles (right). The transparent blue structure is the strand-swap crystal structure (PDB ID: 2QVF). The transparent red structure is the X-dimer crystal structure (PDB ID: 3LNH). The remaining, opaque structures are the simulated structures. **(C)** Root mean squared deviations (RMSD) of the pulled structures relative to the strand-swap dimer crystal structure as a function of the time it took to converge to the strand-swap crystal structure (minimum RMSD). RMSDs were determined by least-squares fitting to the strand-swap dimer crystal structure and calculating the distance between alpha carbons of the two structures. **(D)** Model for vinculin-dependent, force-induced conversion of X-dimers to strand-swap dimers. In the absence of vinculin, force applied is insufficient to convert the X-dimer to a strand-swap dimer (left). Cytoskeletal tension is transmitted in a vinculin dependent manner to the Ecad extracellular bond to promote conversion to a strand-swap dimer (right).

## DISCUSSION

Using live-cell, single molecule AFM measurements, we directly demonstrate inside-out regulation of cadherin conformation and show that cytoplasmic proteins regulate Ecad adhesive properties, similar to inside-out regulation of integrins ^48^. Our data establish a distinct, mechanistic role for vinculin in allosterically regulating Ecad conformation and adhesion at the single molecule level. Our results also demonstrate that previous interpretations of how vinculin strengthens Ecad adhesion are incomplete. While these studies solely focused on vinculin mediated remodeling of the junctional actomyosin cytoskeleton ^22–24^, we demonstrate that vinculin also changes the adhesive properties of individual Ecad ectodomains. Furthermore, our data suggests that vinculin mediates the transmission of cytoskeletal contractile force to the Ecad ectodomain and drives the conversion of X-dimers to strand-swap dimers. Consequently, when Ecads decouple from vinculin and/or α-catenin, they are trapped in an X-dimer structure. Force induced changes in Ecad dimer conformation has been previously suggested ^7^, but until now has not been directly measured.

While previous AFM measurements with α-catenin knockdown cells suggest that reducing α-catenin levels decrease the unbinding force of Ecad ectodomains, the molecular mechanism by which α-catenin influences Ecad adhesion was not determined ^49, 50^. Our data shows that vinculin association and myosin II dependent cytoskeletal contractility allosterically trigger strand-swap dimerization of Ecad and suggests that α-catenin affects Ecad adhesion by serving as the scaffold protein that binds vinculin to Ecad-catenin complex.

It is important to point out that our experiments measure the allosteric conformational regulation of non-junctional Ecads present on the apical surface of cells. While apical Ecads may be organized differently from Ecads localized to cell-cell junctions ^51, 52^, the intracellular proteins that couple Ecad to the actin cytoskeleton are similar for both Ecad pools. Previous studies show that both apical and junctional Ecads are under similar levels of constitutive actomyosin tension ^47^, suggesting that both Ecad pools interact with the actin cytoskeleton in similar manners. Moreover, magnetic twisting cytometry show force-induced, vinculin-dependent stiffening of Ecad coated beads placed on the apical cell surface, suggesting that vinculin also associates with the extra-junctional Ecad ^18^. More recently, 3D super-resolution microscopy of MDCK cells attached to an Ecad-coated substrate, show that the intracellular organization of many Ecad-associating proteins including p-120 catenin, β-catenin, α-catenin and vinculin, resemble native cell-cell junctions ^53^. These studies suggest that the inside-out regulation that we measure with apical Ecads, also occurs with junctional Ecads.

Our ECIS experiments show that while barrier formation half time for both parental-Trp and ΔVBS cells was longer compared to parental cells (Figure 4F), the maximum impedance after barrier formation was completed for parental and ΔVBS cells were similar to each other and slightly higher than parental-Trp cells (Figure S4). This finding is explained by the fact that the maximum impedance is a measure of the steady state of barrier integrity after the dynamic process of junction formation is completed and is principally determined by formation of tight junctions. Consequently, given enough time, the clustering of Ecads within ΔVBS junctions could orient them to promote formation of strand-swap dimers, similar to parental cells. However, when Trp is added to the parental cells, it blocks strand-swapping and traps Ecad in an X-dimer conformation even when junction formation is completed and reaches steady-state. Previous studies show that tight junctions in Ecad deficient mice have increased permeability ^54^. This suggests that inhibiting strand-swapping in Ecad may affect tight junction formation resulting in a lower maximum impedance in parental-Trp cells.

Despite the large number of proteins expressed on the cell surface, we were able to measure single molecule unbinding by limiting Ecad density on the AFM tip and controlling the force with which the AFM tip contacts the cell. However, we were also aided by a low density of Ecads expressed on the apical surface of confluent MDCK cell monolayers. While the precise oligomeric state of Ecad on apical cell surfaces is unclear, with super-resolution imaging suggesting that apical Ecads in A431 cells are present as well separated monomers ^51^ while Ecads in A431D cells organize into small nanoclusters ^52^, our low unbinding event rates nonetheless suggest that only single Ecad on the AFM tip interact with Ecad on the cell surface even if a direct comparison between MDCK and A431/A431D cells cannot be made potentially due to their different properties. Interestingly, the total Ecad event rate measured with αcat-KO cells is higher than with parental, ΔVBS or vinc-KO cells (Figure 2B). This likely occurs because the αcat-KO cells do not form stable cell-cell junctions ^25^. Consequently, Ecads that would otherwise be corralled within cell-cell junctions are now redistributed on the apical cells surface and subsequently interact with Ecads on the AFM tip with a higher probability. The same may be true for other membrane proteins which contributes to an increase in non-specific event rate in control experiments with αcat-KO cells (Figure 2B).

Although AFM is uniquely suited to study the biomechanics of membrane proteins, as with any experiment, there are limitations. The cell surface is a complex environment with many molecules and it is difficult to determine which cell surface protein is interacting with Ecad on the AFM tip. For this reason, we engineered the Ecad-KO cell lines and show that we are not observing any significant heterophilic interactions. For instance, while Ecad can interact heterophilically with several transmembrane proteins, including Desmoglein-2 (Dsg2) ^55, 56^, our control measurements with Ecad-KO cells show that there is no significant binding of Ecad with other cell-surface proteins (Figure 2B). The binding rates measured for the interaction between an AFM tip functionalized with Ecad and the Ecad-KO cells (which corresponds to heterophilic binding between Ecad on the AFM tip and other proteins expressed in MDCK cells) is similar to the binding rate for a bare AFM tip lacking Ecad and parental MDCK cells expressing all transmembrane proteins. This demonstrates that heterophilic Ecad interactions on the apical cell surface occur at such a low rate that we are unable to detect them. One possible reason for the low heterophilic binding of Ecad is that other transmembrane binding partners may have low expression on the apical surface of MDCK cells. Alternatively, the affinity of Ecad heterophilic binding may be lower compared to Ecad-Ecad interactions. In addition, since the structures of Ecad bound to different heterophilic binding partners are unknown, our experimental design may also preclude some of these interactions from occurring. For instance, since Dsg2 binds to the *cis* interface of Ecad ^55^, this would require tethered Ecads on the AFM cantilever to flip their orientation and interact with Dsg2 on the cell surface, an event that has a low probability of occurring. Finally, it is important to note that western blot analysis shows similar level of Dsg2 expression for all cell lines compared to Ecad-KO cells (Figure S5), suggesting that the lack of heterophilic Ecad-Dsg2 binding in Ecad-KO cells would be true for other cell lines as well. However, we cannot completely eliminate the possibility of some heterophilic binding in our experiments as it is possible that the knock-out cells have different localization of Ecad and its binding partners. It is also important to note that since our cells are grown on non-porous glass coverslips for up to 72 hours, they are, at best, only partially polarized. Consequently, while some microvilli or primary cilia may be present on the apical cell surface, the probability of probing Ecad on these structures is quite low.

Previous studies also suggest that α-catenin is not the sole linker of Ecad and the cortical actin cytoskeleton. The binding of β-catenin to vinculin is believed to serve as an alternate, secondary bypass connection between Ecad and F-actin ^57^. However, this alternate coupling is believed to be initiated by α-catenin ^58^. Our AFM data is consistent with these findings. The fraction of cytoskeletal-linked Ecad in both parental and ΔVBS cells, which presumably contain the primary α-catenin/F-actin and the secondary β-catenin/vinculin linkages, are similar (Figure 4A). In contrast, in vinc-KO cells, which presumably contain α-catenin/actin linkages but do not contain β-catenin/vinculin linkages, we see a reduced coupling of Ecad to the actin cytoskeleton (Figure 4A). Finally, in αcat-KO cells we see practically no coupling between Ecad and F-actin, presumably because both the primary α-catenin/actin and the alternate β-catenin/vinculin linkages cannot be formed (Figure 4A). However, our data do not eliminate the possibility that the greater fraction of Ecad-actin linkages in ΔVBS cells compared to vinc-KO cells occurs because the ΔVBS cells contain higher amounts of cytoplasmic α-catenin (Figure 1C).

It is also worth noting that measurements in αcat-KO cells — where Ecad cannot link to actin cytoskeleton — mainly resulted in membrane-tethers (Figure 4A), validating the use of membrane-tethers as a signature of cytoskeleton-uncoupled Ecad. Furthermore, the very low occurrence of jumps and cyto-tethers in αcat-KO cells (Figure 4A) implies that jumps and cyto-tethers indeed correspond to the pulling of Ecad coupled to the underlying cytoskeleton. Additionally, the presence of only jumps in the parental-Trp, vinc-KO, and ΔVBS cells (Figure 4B), where Ecad exclusively formed weaker X-dimers (Figure 5B-C, E), supports that jumps represent X-dimers linked to the actin cytoskeleton. Finally, the decrease in fraction of cyto-tethers (Figure 4B) with the decreased probability of strand-swap dimer formation in parental-Bleb (Figure 5F) strongly suggests that cyto-tethers correspond to strand-swap dimers linked to the actin cytoskeleton. While Ecad in parental cells that are coupled to the cytoskeleton via α-catenin and vinculin (cyto-tethers) exclusively formed strand-swap dimers, we were surprised to observe that Ecad which are decoupled from the cytoskeleton (membrane-tethers) formed X-dimers and strand-swap dimers with coequal probabilities (Figure S3A, B). A likely explanation for this result is that the Ecad-cytoskeletal linkage is dynamic and that even transient interactions with vinculin are sufficient to drive strand-swap dimer formation. Since the AFM-tip and the cell surface remain in contact for about 0.45s in our experiments (Figure S6), this may allow enough time from Ecads on the parental cells to convert to a strand-swap dimer conformation and detach from the cytoskeleton (thereby forming membrane-tethers), before the AFM-tip is withdrawn from the cell surface.

When parental cells were treated with blebbistatin, the reduced actomyosin tension resulted in a decrease in the fraction of cyto-tethers (Figure 4B). While these cyto-tethers behaved like parental cyto-tethers and exclusively formed slip-bonds (Figure S3C), a larger fraction of parental-Bleb membrane-tethers formed X-dimers compared to parental cells (69% in parental-Bleb compared to 55% in parental cells), further demonstrating that myosin generated contractile force is important for cadherin strand-swap dimer formation (Figure S3D).

Given the higher lifetimes for X-dimers compared to strand-swap dimers at forces > 50 pN (Figure 5A), it might be expected that force would convert Ecad to X-dimers instead of strand-swap dimers. However, since contractile force applied by actomyosin on a single Ecad is only around 1-2 pN ^47^ and the lifetime of the strand swap dimer is higher than the X-dimer at forces < ∼50 pN, (Figure 5A), the strand-swap dimer is the preferred binding conformation in a physiological context. Furthermore, as shown by our simulations (Figure 6A-C), the geometry of these conformations do not permit a transition from strand-swap dimer to X-dimer since pulling on the strand-swap dimer moves the X-dimer interface further apart.

Consistent with a previous study that used optical tweezers and single particle tracking to infer the fraction of cytoskeleton-bound Ecad ^36^, our data shows that only ∼50% of Ecads on the apical cell surface are coupled to the actin cytoskeleton. Studies also show that cells expressing Ecad/α-catenin fusion constructs that constitutively bind to the actin cytoskeleton, exhibit higher cell-cell adhesion and reduced intercellular motility compared to cells expressing wild type Ecad ^36, 59^. These findings are supported by our observation that two rapidly exchangeable pools of Ecad structures (X-dimers and strand-swap dimers) exist on the cell surface. It is likely that these different pools of Ecads are crucial in reorganizing cell-cell junctions during collective cell migration: as the cell-cell contacts rapidly rearrange, strand-swap dimers anchor the contact while X-dimers probe for formation of new contacts.

Based on the results presented in this manuscript, we propose a biophysical model for the inside-out regulation of Ecad conformation. Our data suggests that vinculin mediated myosin II recruitment generates a contractile force which propagates to the Ecad ectodomain and drives the conversion of X-dimers to strand-swap dimers. (Figure 6D). Consequently, treating parental cells with blebbistatin, inhibits myosin II dependent cytoskeletal contractility and reduces the ability of Ecad to form robust strand-swap dimers (Figure 5F). All atom computer simulations also demonstrate that a force applied to the membrane proximal region of X-dimers results in formation of strand-swap dimers (Figure 6B, 6C). Previous studies suggest that Ecad adhesion is altered by changing the phosphorylation state of cytoplasmic p120-catenin ^40, 60, 61^; however, the mechanism by which this occurs and the exact conformational states Ecads adopt, are unknown. It is possible that p120-catenin also indirectly modulates vinculin association with the Ecad cytoplasmic region, which in turn regulates Ecad ectodomain conformation and adhesion.

Recent studies have suggested that regulation of Ecad adhesion plays an important role in cancer progression. For instance, it has been shown that up-regulation of Ecad adhesive activity, rather than its amount, reduces the number of cells metastasized from the mammary gland to the lung ^62^. Additionally, several point mutations in Ecad ectodomains that are prevalent in hereditary diffuse gastric carcinoma selectively interfere with the inside-out regulation of Ecad adhesion, rather than the ability of Ecad to adhere ^62^. Furthermore, studies have shown that while vinculin expression is significantly reduced in colorectal cancer (CRC), overexpressing vinculin reduces metastasis in CRC ^63^ and other cancer cell lines ^64^. Consequently, understanding how cytoplasmic proteins induce conformational changes of Ecad ectodomains could be crucial to understanding the role of Ecad in cancer progression and metastasis.

## METHODS

### Generating mutant cell lines

Generation of αcat-KO cells (ΔαE-catenin MDCK cells) ^25^ has been described previously. For generation of ΔVBS cells, αcat-KO cells were stably rescued ^26^ with GFP-α-catenin–ΔVBS mutants ^35^ as described previously. Vinc-KO and Ecad-KO cells were generated by stably expressing a PiggyBac plasmid encoding Cas9 with a cumate inducible promoter, and gRNA expression cassette for canine Ecad or vinculin specific gRNA, which resulted in complete knockout cells (see Figure 1C). The mutations were sequence verified by PCR amplifying the target sequences from genomic DNA of knockout cell lines and inserted into TOPO vectors (Invitrogen) for sequencing.

Ecad:

gRNA: ACAGACCAGTAACTAACGA

KO: ACAGACCAGT----AACGA

Vinculin:

gRNA: GGA GCACCGAGTAATGTTGG

KO: GGAtGCACCGAGTgATGTTGG

### Western Blotting

Cell lysates for western blotting were prepared in 1x SDS sample buffer (10 mM Tris pH 6.8, 1% SDS, 10% glycerol, 0.005% Bromophenol Blue, 2% β-mercaptoethanol). Primary antibodies against E-cadherin (clone 36; BD Biosciences), α-catenin (clone 15D9; Enzo Life Sciences), vinculin (clone hVIN-1; Milipore Sigma), Dsg2 (clone 141409; R&D Systems), and α-tubulin (clone DM1A; Cell Signaling) were used in manufacturer recommended dilutions in Phosphate-Buffered Saline with 0.1% Tween 20 (PBST). Horse Radish Peroxidase (HRP) conjugated secondary antibody (catalog no. 1706516; Bio-Rad) was used in 1:1000 dilution in PBST. Protein samples were detected with a WesternBright Chemiluminescence Kit (catalog no. K-12045; Advansta). Images were acquired using Image Lab software from Bio-Rad.

### Purification of Ecad ectodomain

Generation of Ecad monomer plasmids containing a C-terminal Avi tag has been described previously ^65^. The plasmids were incorporated into pcDNA3.1(+) vectors and were transiently transfected into HEK 293T cells using PEI (Milipore Sigma) as previously described ^66^. Three days post transfection, conditioned media was collected for protein purification. Purification of Ecad were performed using methods described previously ^8, 65^. Media containing his-tagged Ecads was passed through a chromatography column containing Ni-NTA agarose beads (Qiagen). Beads were then washed with a pH 7.5 biotinylation buffer (25mM HEPES, 5mM NaCl, and 1mM CaCl_2_). Ecads bound to the Ni-NTA beads were biotinylated with BirA enzyme (BirA 500 kit; Avidity) for 1hr at 30^0^C. Following biotinylation, free biotins were removed using the Ni-NTA column and biotinylated Ecads bound to Ni-NTA beads were eluted using a Ph buffer containing 200mM Imidazole, 20mM Na_2_HPO_4_, 500mM NaCl, and 1mM CaCl_2_.

### Epithelial barrier formation measurements

Ecad-dependent formation of epithelial barriers was assessed by real-time impedance sensing. To this end, parental or ΔVBS cells were trypsinized, re-suspended in calcium-free DMEM medium (Immunosource, D9800-10) supplemented with 10% dialyzed fetal calf serum, and plated onto L-cysteine-reduced, collagen I-coated 8WE10 electrodes (Applied Biophysics) at a density of 2×10^5^ cells per well and in the presence or absence of 2 mM Trp. Impedance measurements were performed at 37°C and 6% CO2 using a 1600R Electrical Cell Impedance Sensing (ECIS) system (Applied Biophysics) at a frequency of 4000 Hz. CaCl_2_ was added at a final concentration of 2 mM to allow for adherens junction formation, and measurements were continued until stable impedance levels were reached in all conditions. Within each individual experiment, 4 technical replicates for each condition were averaged to assess the time-dependent impedance for that condition. The half times of barrier formation were calculated as the timepoint at which the impedance signal surpassed 50% of the ultimate maximum impedance.

### Ecad functionalization on AFM cantilever

Purified Ecad monomers were immobilized on Si tip of AFM cantilevers (Hydra 2R-100N; AppNano) as described previously ^8, 9, 55, 65^. Briefly, the AFM cantilevers were cleaned by immersing in 25% H_2_O_2_/ 75% H_2_SO_4_ solution overnight. The cantilevers were then sequentially washed with deionized water and acetone. The cantilevers were functionalized with amine groups by immersing in 2% (vol/vol) 3-aminopropyltriethoxysilane (Millipore Sigma) solution dissolved in acetone. Polyethylene glycol (PEG) (MW 5000, Lysan Bio) spacers containing amine-reactive N-hydroxysuccinimide ester group at one end were covalently attached to the cantilever (100 mg/ml in 100mM NaHCO_3_ dissolved in 600mM K_2_SO_4_, for 4 hrs); 20% of the PEG spacers presented biotin molecules at the other end. Unreacted amine groups on the silane molecules were capped using NHS-sulfo acetate (10 mg/ml for 30 mins; Thermo Fisher). PEG-functionalized AFM cantilevers were incubated in 1 mg/ml Bovine Serum Albumin ^55^ overnight. The cantilevers were then sequentially incubated with streptavidin (0.1 mg/ml for 30 mins) and biotinylated Ecads (200 nM for 90 mins). Finally, free biotin-binding sites of streptavidin were blocked by using 0.02 mg/ml free biotin for 20 mins. All steps, except the incubation with Ecad, were kept the same for cantilever functionalization in control experiments.

### Cell culture for AFM experiment

Cells were cultured in low glucose Dulbecco’s modified Eagle’s medium (DMEM) (Milipore Sigma) supplemented with 10% FBS (Milipore Sigma) and 1% antibiotics solution (penicillin + streptomycin + kanamycin) at 37^0^C and 5% CO_2_. Cells were trypsinized 48-72 hrs prior to the experiment and plated on an ethanol-cleaned glass coverslip at a low density. The confluent monolayer of cells was washed 5 times with a pH 7 DMEM and antibiotics solution before loading onto the AFM setup with the same medium.

### AFM force measurements

Force measurements were performed using two Agilent 5500 AFMs with closed loop scanners. An AFM cantilever functionalized with purified Ecad was brought in contact with a confluent monolayer of cells, pressed against the cell surface with a low force (typical pressing force was about 50 pN) for 0.1s, and then retracted at one of five constant velocities (3, 5, 7, 9, 11 µm/s). Control experiments used cantilevers decorated with streptavidin but lacking Ecad. Cantilever spring constants were measured using the thermal fluctuation method ^67^. All the experiments were performed in pH 7 DMEM and antibiotics solution in a custom-built environmental chamber supplied with 5% CO_2_ at room temperature. Parental-Bleb and parental-Trp experiments were performed with 100μM (±) -Blebbistatin (Milipore Sigma) and 2mM L-Tryptophan (Alfa Aesar) in the solution respectively. A typical experiment lasted for ∼10 hours yielding ∼1200 force-distance traces.

### Analysis of AFM data

AFM-generated force-distance traces were analyzed using custom MATLAB scripts. Traces containing single rupture events were classified into membrane-tether, jump, and cyto-tether categories based on the following criteria:

- Membrane-tethers: force plateau at least 100nm long and slope less than 25 pN/μm.
- Cyto-tethers: have a difference between the peak force and the plateau force that is greater than the noise, calculated as 2 standard deviations of 500 data points after the force drop.
- Jumps: have a linear force-distance relationship with a slope of more than 25 pN/μm.

Rupture events having unbinding force less than the noise of each respective force curve or a force >150 pN were not considered for analysis. Tethers were further fit to the SLS model as described in ^31^ while jumps were fit to a Hookean spring model. We only retained events that fit either the SLS or spring model with a Root Mean Squared Error (RMSE) of less than the mean plus two standard deviations of all fits for each respective model.

To obtain the fractions of membrane-tether, cyto-tether, and jump events for each cell line, we first measured the binding probability of each type of event for both the Ecad experiment and the control. The binding probabilities obtained from the control experiment were subtracted from the corresponding event rate of the Ecad experiment.

Loading rates for the jump events used for dynamic force spectroscopy analysis (Figure 4C-E) were calculated by multiplying the slope of the jump by the pulling velocity. K-means clustering was used to group the loading rates as described previously ^68^. Mean unbinding force and mean loading rate for the groups were fit using a weighted non-linear least square fit to the Bell-Evans model ^37^. Raw data was bootstrapped with replacement, grouped using k-means, and the mean values were fit to the Bell-Evans model 1000 times; the fitting parameters and errors were determined from the mean and standard deviations of the resulting distributions. The data used in each fit was weighted by the fraction of data points in a particular bin relative to the total number of data points. Force-dependent Ecad-Ecad bond mechanics were studied by analyzing the forces and lifetimes of membrane-tethers and cyto-tethers. Bond lifetimes were determined from the force curves starting at the beginning of the plateau region to the bond rupture. The unbinding force was calculated as the mean of the force values spanning that same region. This data was used to create the plots shown in Figure 5. The unbinding forces were fit to a Gaussian mixture model (GMM) using an expectation maximization algorithm to determine the bins; the means and standard deviations used in the GMM fit were initialized using the k-means clustering algorithm. We determined the characteristic lifetime of each bin using maximum likelihood estimation (MLE) on bootstrapped samples. The likelihood function, 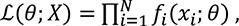 was formulated using an exponential distribution *f*.

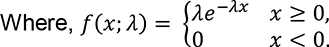

The MLE lifetimes were fit by Chi-squared minimization to one of two models: a composite slip-catch bond model or a catch bond model ^46^. Assuming the errors in the measurements are Gaussian distributed, fitting by minimizing Chi-squared is the same as maximizing the log likelihood ^46^. The composite slip-catch model used to fit the parental and parental-Bleb data was the sum of a slip bond model and a catch bond model. The slip bond model ^45^, 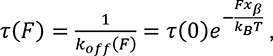 was used as the first component of the composite slip-catch model to fit the parental and parental-Bleb data (Figure 5A, F), where *τ* is the characteristic bond lifetime in the absence of force and *x*_*β*_ is the reaction coordinate distance. The catch bond model used in the composite model and to fit all other cell lines is given by,

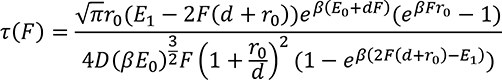

where *r*_0_ is the natural bond length, *d* is the reaction coordinate (same as *x*_*β*_ for the slip bond model), *E*_0_ and *E*_1_ are components of the bond energy, 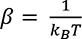 is the Boltzmann constant), and *D* is a diffusion constant 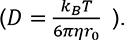. The number of bins used was established by the minimum weighted RMSE of the composite slip-catch or catch bond model relative to the mle lifetimes when fitting the forces with the GMM using 5 to 10 bins. (7-10 for parental and parental-bleb) for each cell line. The weights were determined by the fraction of data points in a bin relative to the total number of data points for that cell line. The composite slip-catch and catch bond model fitted parameters are shown in Table S2. The Chi-squared fitting function was acquired from the Mathworks File Exchange website (Generalized Nonlinear Non-analytic Chi-Square Fitting by Nathaniel Brahms, May 26, 2006).

The fraction of strand-swap and X-dimers calculated from Figure 5A and 5F were determined by integrating each component of the composite model in 5 pN bins from 0 to 150 pN to determine the slip/catch area of that bin. The number of data points in each bin that corresponded to the slip bond model, for example, was determined by,

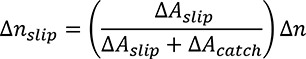

where Δ*A*_*slip*_ and Δ*A*_*catc*ℎ_ are the area under slip and catch bond fit respectively for the 5pN bin, and Δ*n* is the total number of data points in the 5pN bin. As expected, the sum of the slip and catch bond fits—and therefore the sum of their areas—is equal to the composite fit. The total fraction of data corresponding to the slip bond (strand-swap dimers) is then,

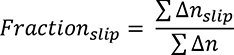

### Steered Molecular Dynamics Simulations and Structural Analysis

Steered molecular dynamics simulations were performed with GROMACS 2020.1 using the FARM high performance computing cluster at UC Davis. The X-dimer crystal structure (PDB ID: 3LNH) was used in all simulations. Missing residues were added using PDBFixer and the N-terminal β-strand with Trp2 was positioned near the strand-swap interface using the sculpting tool in PyMOL. All simulations were performed with the OPLS-AA/L force field and the TIP4P water model.

The X-dimer crystal structure was equilibrated by performing a 20 ns molecular dynamics (MD) simulation. To setup the simulation, the X-dimer was placed in the center of a dodecahedral box such that no atom of the X-dimer was closer than 1 nm to any boundary. The box was solvated by adding water molecules and charge-neutralized by adding Na^+^ ions. Additionally, NaCl, KCl, and CaCl_2_ were added to the box at concentrations of 150 mM, 4 mM, and 2 mM, respectively, for a total system size of 148,965 atoms. Energy minimization, using the steepest decent algorithm, was performed until the force on any atom in the system was less than 1000 kJ/mol/nm. The temperature, pressure, and density of the system were stabilized by equilibrating under *isothermal-isochoric* and *isothermal-isobaric* conditions using a modified Berendsen thermostat and Berendsen barostat. Following equilibration, a 20 ns MD simulation was performed with 2 fs integration steps, LINCS constraints on bond length, and long-range interactions were evaluated using the particle mesh Ewald method. The stability of the X-dimer was monitored by calculating the root mean squared deviation (RMSD) of the X-dimer relative to the initial structure.

Conformational switching was modeled using Steered molecular dynamics (SMD) simulations performed on the fully equilibrated X-dimer by pulling on a group of residues (residues 151-166, 174-186, and 208-213) — to avoid unfolding—at the C-terminal end of one Ecad monomer in five directions (0°, ±30°, ±40°; *see Figure 6A*). The C-terminal of the opposing Ecad was constrained to simulate a robust linkage to the actin cytoskeleton. The X-dimer structure from the final frame of the MD simulation was placed at the center of a 12 x 30 x 8 nm^3^ box, and solvated and equilibrated as described above to an average of 376,743 atoms. After centering the X-dimer in the box, the pulling angle was achieved by rotating the X-dimer about the z-axis (short axis) by the corresponding pulling angle. For each simulation, the X-dimer was pulled with a constant force (force constant = 498.2 pN/nm; 300 kJ/mol/nm^2^) along the long axis of the box.

The conversion of the X-dimer to strand-swap dimer was determined by aligning the simulated structures to the X-dimer (PDB ID: 3LNH) and strand-swap dimer (PDB ID: 2QVF) crystal structures and calculating the RMSD of the alpha carbon atoms before and after pulling, respectively. The strand-swap dimer was formed from 2QVF by crystallographic symmetry operations.

## Acknowledgments

Research in the SS lab was supported in part by the National Science Foundation (PHY-1607550) and National Institute of General Medical Sciences of the National Institutes of Health (R01GM121885 and R01GM133880). Research in the SY lab was supported by NIH R03 EB021636, NSF 1562095 (SY, REU supplement to JC), and UC Davis Provost’s Undergraduate Fellowship (to JC). JC is a recipient of the Beckman Scholars Award. Research in the MG lab was supported by the Netherlands Organization for Scientific Research (NWO; 016.Vidi.189.166 and NWO gravitational program CancerGenomiCs.nl).

## Data availability

All data produced by this research are made available in the Supporting Information.

## Author contributions

R.K., A.V.P. and S.S. designed research; S.S. directed the research; R.K., A.V.P. and C.F.Y. performed AFM experiments; A.V.P. performed computer simulations; W.J.P and M.G. performed ECIS experiments; J.S.C., M.G., and S.Y., generated cell lines; R.K., A.V.P., and S.S. wrote the paper.

## Notes

### Competing Interest Statement

The authors have declared no competing interest.

