## Supplemental Information for "Inside-out regulation of E-cadherin conformation and adhesion"

[#] Equal contribution

**Note 1: Standard linear solid (SLS) model used to fit membrane-tethers and cyto-tethers.**

This viscoelastic model consists of a spring with stiffness, $k_t$, that is connected in parallel to another spring ($k_c$) and dashpot ($\mu_c$) in series.

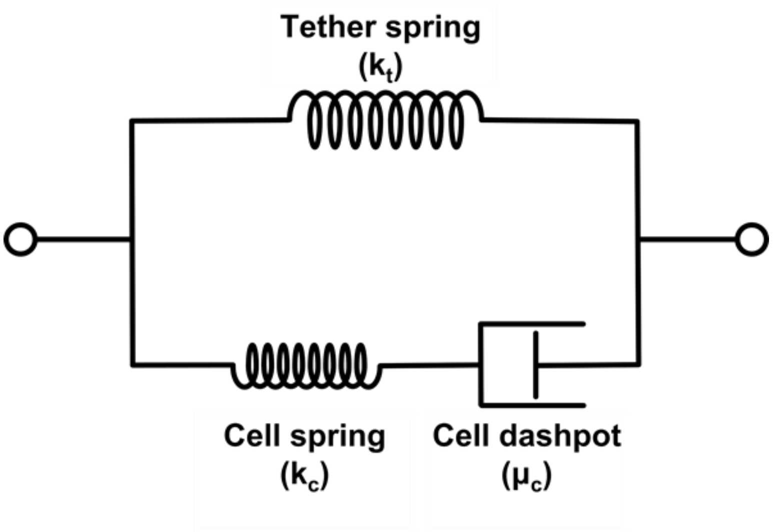


As described in ref. 30, the stiffness of the tether is described by $k_t$, while the cell spring, $k_c$, corresponds to the cell membrane rigidity and $\mu_c$ describes the cell viscosity. The differential equation for the SLS model is

$$\dot{F}(t) = -\frac{k_c}{\mu_c}(F(t) - k_t v t \; - \mu_c v(1 + \frac{k_t}{k_c}))$$

Solving this with the initial condition $F(t_0) = F_0$ to accurately model the nonzero initial force for cyto-tethers gives

$$F(x) = (F_0 \; + \; k_t \, x_0 \; - \mathrm{v}\mu_c) \, e^{-\frac{k_c(x - x_0)}{v \, \mu_c}} + \; k_t \, x + \mathrm{v}\mu_c$$

Where we have used $x = v \cdot t$ for pulling at a constant velocity.

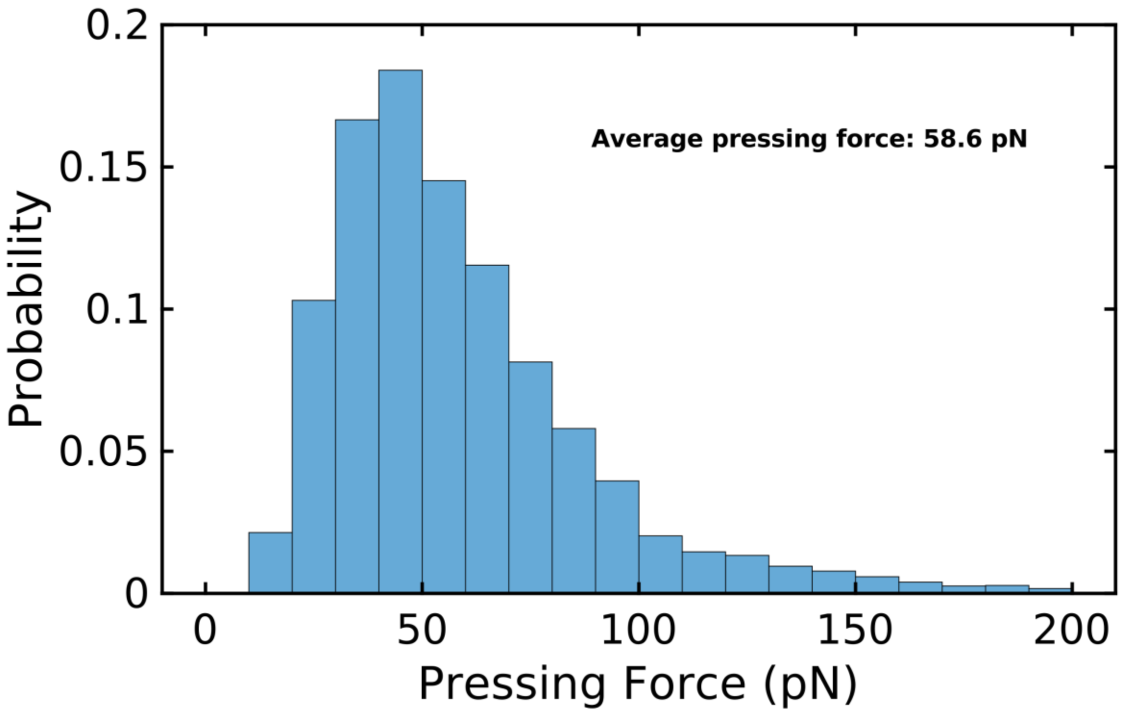


**Figure S1: Histogram of the initial pressing force on MDCK cells in AFM experiments.** The mean pressing force is 58.6 pN as calculated from the retract traces.

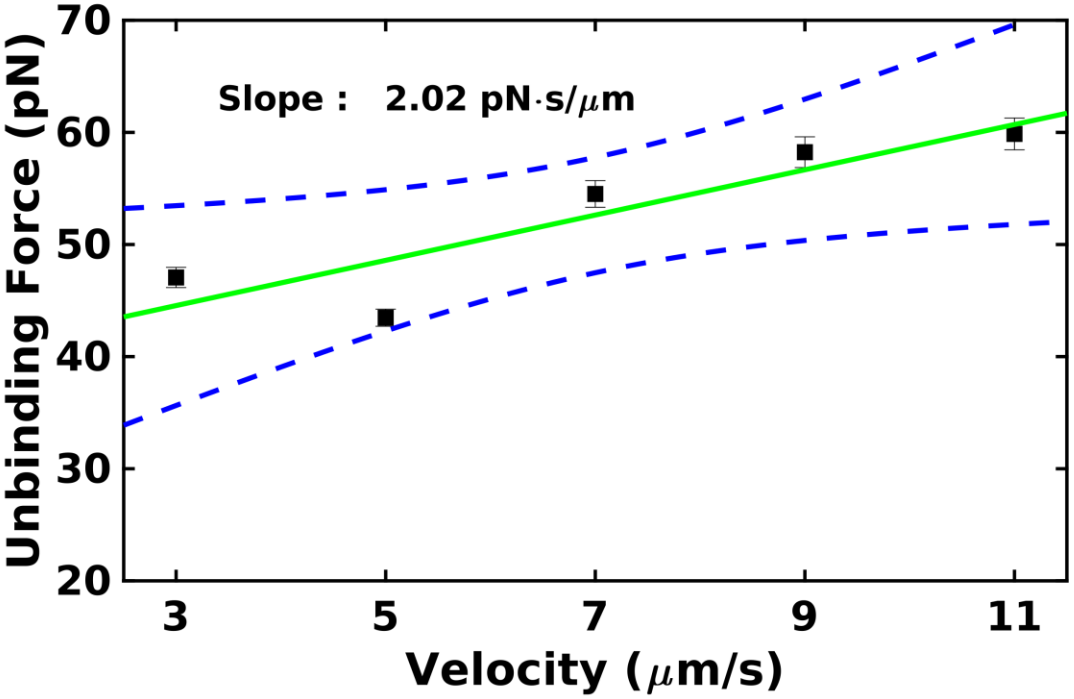


**Figure S2: Tether unbinding force linearly correlates with pulling velocity.** Slope = 2.02 pN/(µm/s). Dashed blue lines represent 95% confidence bounds of the fit. Errors are the standard deviations of the bootstrapped means.

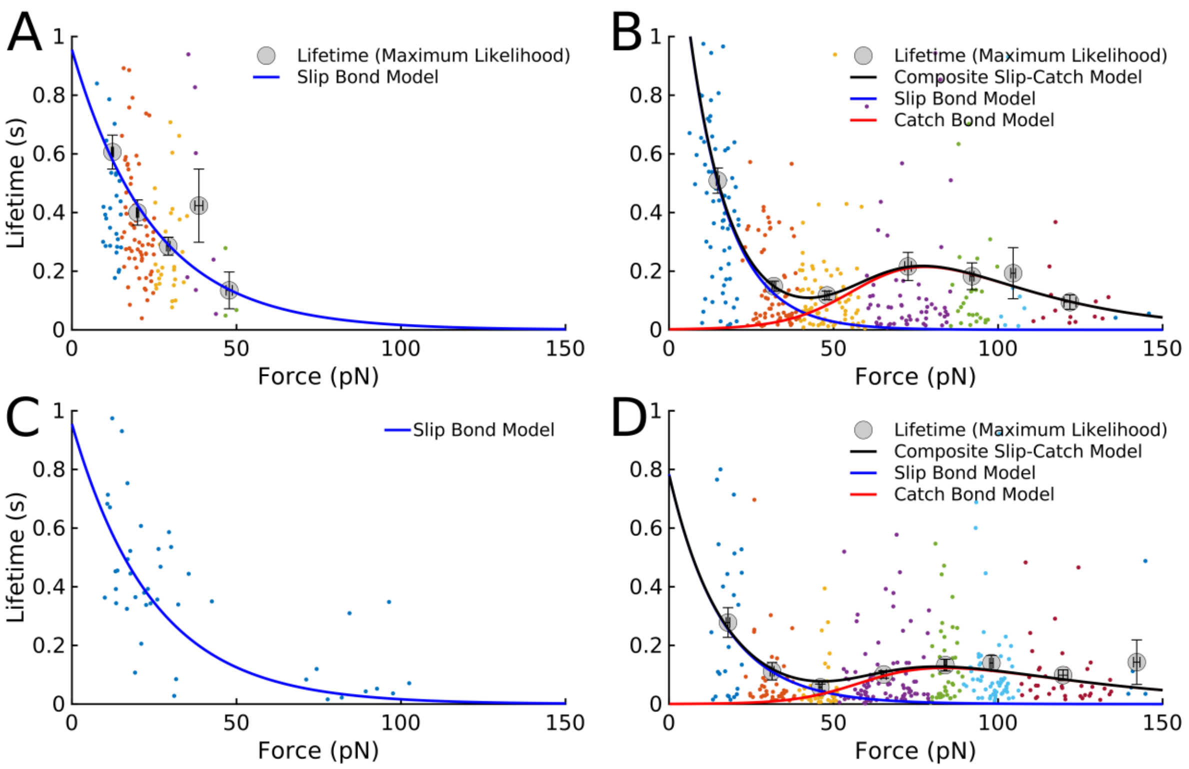


**Figure S3: Force vs lifetime for cyto-tether and membrane-tethers in parental and parental-bleb cells. (A)** In parental cells, all cyto-tethers exhibit slip bond behavior while **(B)** membrane-tethers form both catch bonds (55%) and slip bonds (45%). **(C)** Cyto-tethers in parental-bleb cells also form slip bonds while **(D)** membrane tethers form slip and catch bonds albeit at a reduced slip bond fraction (31% slip bonds, 69% catch bond). This is presumably the result of reduced actomyosin contractility in parental-bleb cells. Due to limited number of data points, the cyto-tether data in (A) was binned using a fixed bin width (10 pN) instead of binning using the Gaussian mixture model. The membrane-tether data in (B) and (D) were binned using the Gaussian mixture model (as in Figure 5) and fit to the composite slip-catch model. The slip bond fit for parental cyto-tethers (A) was overlaid to the parental-bleb raw data (C) since proper binning and fitting was not possible considering the sparse cyto-tethers for parental-bleb cells.

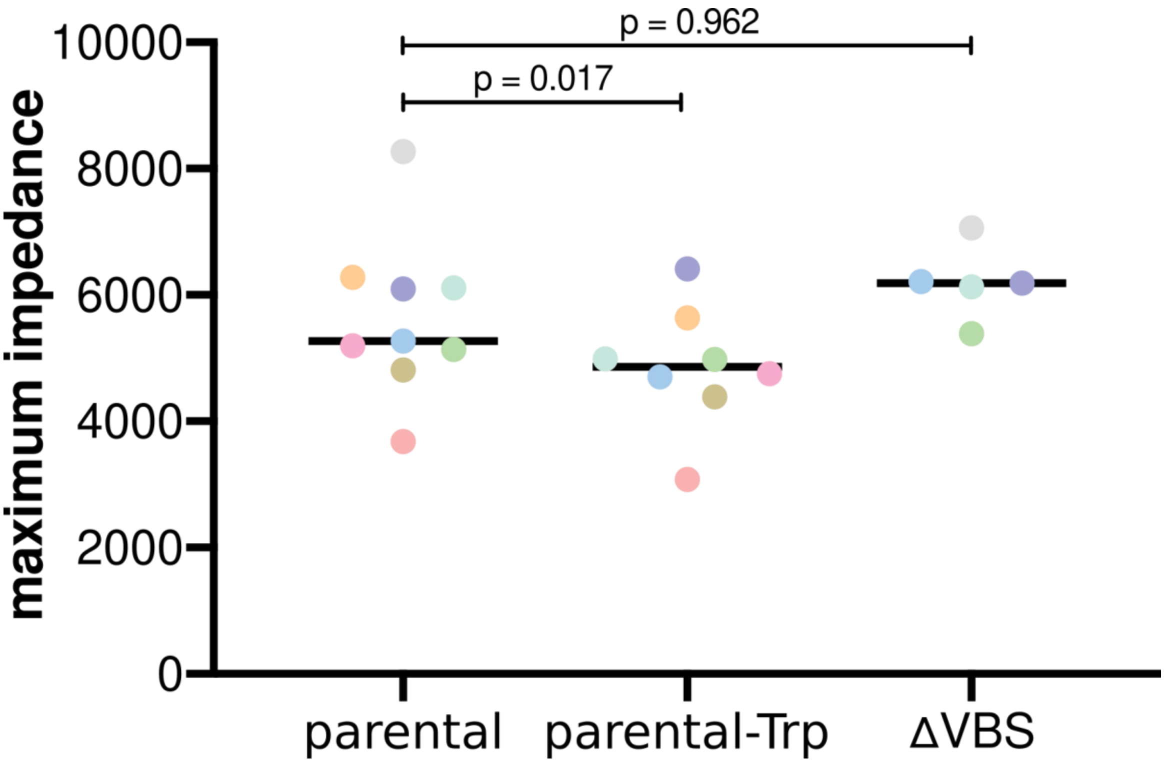


**Figure S4: Trans-epithelial resistance maximum impedance.** Trans-epithelial resistance measurement showing maximum level of impedance following calcium-dependent formation of an epithelial barrier in parental cells in the absence (n = 9) and presence (n = 8) of 2 mM tryptophan, or ΔVBS cells (n = 5). Colors indicate conditions that were measured within the same experiment. Paired t-test.

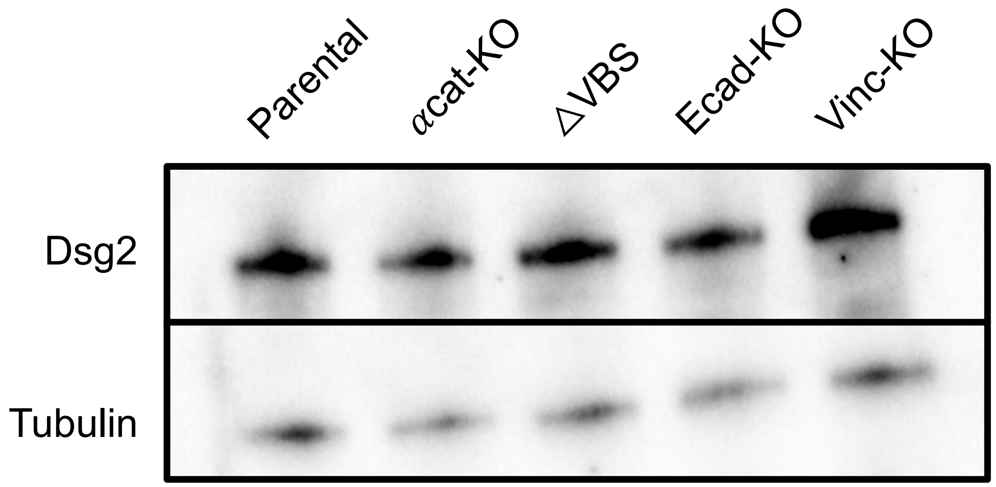


**Figure S5: Western blot analysis of cell lysates for Dsg2.** Mutant cells express similar level of Dsg2 compared to parental cells (αcat-KO – 105%, ΔVBS – 109%, Ecad-KO – 94%, and Vinc-KO – 120%). Bands were quantified using Image Lab software from Bio-Rad.

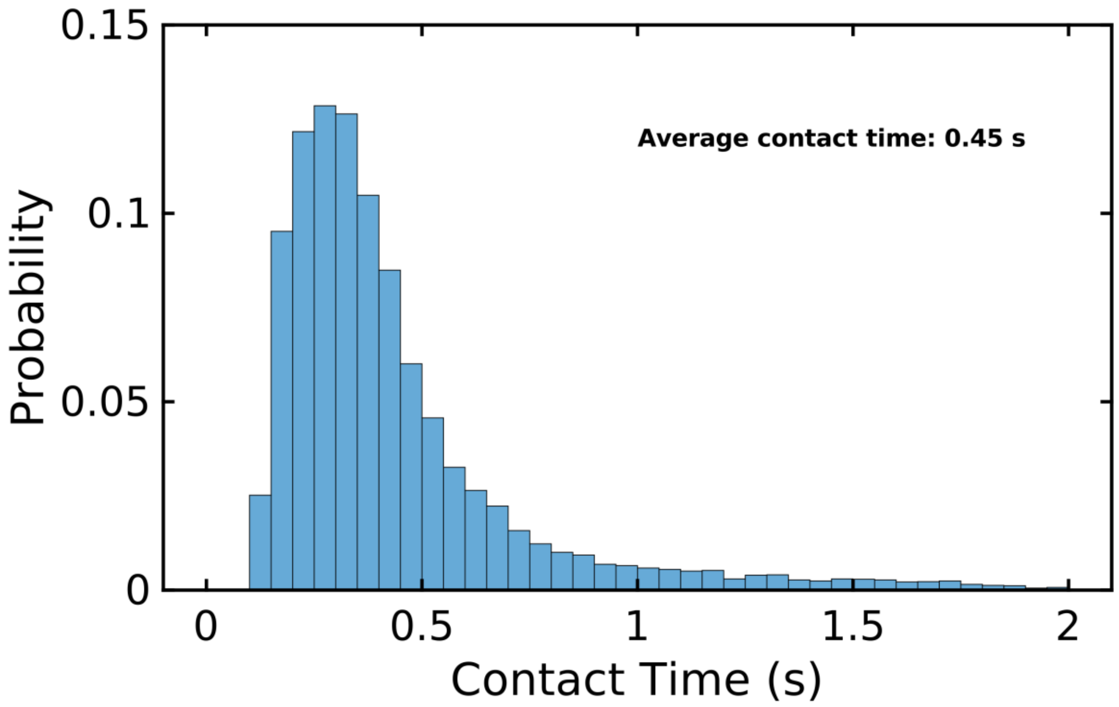


**Figure S6: Histogram of tip surface contact duration.** AFM tip remained in contact with the cell surface for 0.45s on average before pulling on the Ecad-Ecad bond. The x-axis was limited to 2 s for clarity.

**A**

| | Median of cyto-tether fit parameters | Median of membrane tether fit parameters |
|---|---|---|
| $k_c$ (μN/m) | 16 ± 2.0 | 543 ± 10 |
| $\mu_c$ (μN·s/m) | 1.8 ± 0.18 | 8.1 ± 0.13 |
| $k_t$ (μN/m) | 1.4 ± 0.26 | 1.0 ± 0.19 |

**B**

| | Median of membrane tether fit parameters | Integrins on Jurkat cells (ref. 31) | | P-selectin on T-cells (data from ref. 32, fit parameters calculated in ref. 31) | |
|---|---|---|---|---|---|
| | | Resting integrins | Activated integrins | -Latrunculin | +Latrunculin |
| $k_c$ (μN/m) | 543 ± 10 | 260 | 190 | 190 | 51 |
| $\mu_c$ (μN·s/m) | 8.1 ± 0.13 | 5.9 | 6 | 9.9 | 4.6 |
| $k_t$ (μN/m) | 1.0 ± 0.19 | 1.6 | 0.9 | 8 | 3 |

**Table S1: SLS fitting parameter values. (A)** Comparison of SLS fitting parameters for cyto-tethers and membrane-tethers. The tether stiffness, $k_t$, is similar for cyto-tethers and membrane-tethers, but cell stiffness ($k_c$) and cell viscosity ($\mu_c$) are less for cyto-tethers. Tether stiffness is expected to be the same in cyto-tethers and membrane-tethers since in both cases the tether is formed without linkage to the cytoskeleton. In contrast, $k_c$ and $\mu_c$ may be different for different initial conditions – *i.e.* tether relaxing from an initial force (cyto-tethers) or pulling the membrane from rest (membrane-tethers). An additional factor may be differences in the cytoplasmic composition within the tether itself since membrane-tethers may contain high densities of cytoplasmic proteins which initially increase resistance to pulling, whereas cyto-tethers , which form after cytoskeletal decoupling, may be able flow more freely yielding lower values for both $k_c$ and $\mu_c$. **(B)** Comparison of SLS parameters for membrane-tethers in this work and in previous studies with integrins on Jurkat cells (ref. 31) and P-selectin on T-cells (data from ref. 32; parameters calculated in ref. 31). Errors were not reported in those studies. Errors for this work were acquired from the bootstrapped medians.

| A | τ (s) | $x_\beta$ (nm) | $r_0$ (nm) | d (nm) | $E_0$ ($k_BT$) | $E_1$ ($k_BT$) |
|---|---|---|---|---|---|---|
| Parental | 1.1 ± 0.23 | 0.22 ± 0.05 | 0.11 ± 0.17 | 0.24 ± 0.04 | 22.3 ± 4.5 | 13.6 ± 3.1 |
| Parental-Bleb | 1.1 ± 0.51 | 0.26 ± 0.08 | 0.50 ± 0.12 | 0.10 ± 0.04 | 12.5 ± 3.5 | 17.4 ± 2.6 |

| B | | | $r_0$ (nm) | d (nm) | $E_0$ ($k_BT$) | $E_1$ ($k_BT$) |
|---|---|---|---|---|---|---|
| Parental-Trp | - | - | 0.10 ± 0.33 | 0.28 ± 0.06 | 24.3 ± 3.0 | 11.5 ± 1.7 |
| Vinc-KO | - | - | 0.23 ± 0.97 | 0.16 ± 0.08 | 27.1 ± 6.1 | 3.8 ± 5.2 |
| αcat-KO | - | - | 0.27 ± 0.29 | 0.10 ± 0.04 | 27.0 ± 2.8 | 3.9 ± 2.0 |
| ΔVBS | - | - | 0.14 ± 0.10 | 0.13 ± 0.05 | 25.3 ± 1.4 | 7.4 ± 1.0 |

**Table S2: Slip and catch bond parameters. (A)** The fitting parameters obtained from the composite model (Figure 5A, F). Parameters for slip bonds are consistent with previous studies of classical cadherins (ref. 8, ref. 38-41) while parameters for catch bond are comparable to the values in *(B)*. **(B)** Catch bond fitting parameters for parental-Trp, vinc-KO, αcat-KO, and ΔVBS (Figure 5B-E). Errors in (A) and (B) were obtained by bootstrapping.
